# Oxidative stress, telomere length, and frailty in an old age population

**DOI:** 10.1101/414680

**Authors:** José Darío Martínez-Ezquerro, Aleida Rodriguez-Castañeda, Mauricio Ortiz-Ramirez, Sergio Sanchez-Garcia, Haydee Rosas-Vargas, Rosalinda Sanchez-Arenas, Paola Garcia-delaTorre

## Abstract

**Background:** A global aging population requires focusing on the risk factors for unhealthy aging, preventive medicine, and chronic disease management. The identification of adverse health outcomes in older adults has been addressed by the characterization of frailty as a biological syndrome. On the other hand, oxidative stress and telomere length have been suggested as biomarkers of aging.

**Objective:** To study the association of oxidative stress, telomere length, and frailty in an old age population.

**Methods:** This was a cross-sectional study based on 2015 data from 202 members from a cohort of older adults (n=202; gender F/M ratio: 133/69; mean age: 69.89 ± 7.39 years). Reactive oxygen species (ROS) were measured by dichlorofluorescin diacetate, and lipid peroxidation by malondialdehyde. Telomere length was determined using qPCR with SYBR Green Master Mix.

**Results:** Statistical analysis showed an association between telomere length and frailty but no association between oxidative stress on telomere length or frailty.

**Conclusions:** Telomere length could eventually be used as a marker to discriminate between healthy and unhealthy aging as expressed by frailty phenotype. However, oxidative stress seems as just a biological process of aging.

## Introduction

A global aging population has motivated researchers to focus on risk factors, preventive medicine, and chronic diseases in order to prepare for the coming health issues that this phenomenon will cause. Economy and society itself will suffer from the change in population age distribution around the globe. Between the years 2015 and 2050 the proportion of the world’s population with more than 60 years of age (1) will increase from 900 million to 2000 million, an increase from 12 to 22% (1). In 2015, Mexico had one older adult (60 years of age and older) for every 10 young adults (15 years old), a figure that is expected to double by 2050 (2). At present, Mexico City is the entity with the highest index of aging in Mexico with 59.1 in 2014 and a projection of 108.7 by 2030 (3). Due to this global demographic transition, successful aging (avoidance of disease/disability, high physical and cognitive function, and sustained engagement with social activities) has become a priority (4).

The concept of aging suggests the emergence and accumulation of chronic conditions such as heart disease, diabetes mellitus (DM), cancer, and cerebrovascular diseases, which are also the main causes of death at this stage of life. Hence, the identification of adverse health outcomes in older adults has been addressed by the characterization of frailty as a biological syndrome. Frailty has been described as the loss of physiologic reserve and resistance to stressors that involves multiple physical, mental, and emotional deficits (5). As frailty advances, vulnerability to dependence increases, and the risk of adverse events such as functional decline, falls, hospitalization, institutionalization, and mortality becomes more plausible (5, 6). In Mexico City, a prevalence of frailty of 20.6% among older adults has been reported (7), an intermediate level when compared with global reports from 4% to 59.1% (8).

Hence the search for biomarkers that can help define frailty at a biological level could be used to improve the diagnosis of frailty, to identify pre-frail patients at an earlier stage, and to evaluate the biological outcome when frailty is modified by factors such as healthy eating or physical exercise. On the matter, telomere length, which has been associated with lifespan (9), has had a controversial role in frailty description. Although no correlation was found for Chinese (10) and German (11) populations, we recently observed shorter telomeres associated with frailty in a Mexican population (12).

Telomeres are protective and regulatory structures of the genome, their shortening is associated with chronic degenerative diseases such as diabetes (13), obesity, and hypertension (14, 15), and has been associated with lifespan (16). Lifespan depends on many internal and external factors such as aging, pollution, quality of life, and education, among others. Consequently, telomere shortening and frailty would seem to be intimately related.

Most of the associations with telomere length mentioned above involve the production of oxidative stress that can affect DNA in a direct form. Oxidative stress has been associated with chronic damage in several conditions including DM and hypertension (9, 17–19). At a molecular level, oxidative stress is capable of altering DNA and has been linked to telomere length reduction (20). In order to determine the possible association between reactive oxygen species (ROS), telomere length and frailty, we evaluated the effect of oxidative stress on telomere length as a first approach to explain telomere length shortening in frail patients. Then, we assessed both telomere length and oxidative stress, as frailty biomarkers.

## Materials and methods

### Participants

This is a cross-sectional study based on the baseline 2015 data from the “Cohort of Obesity, Sarcopenia, and Frailty of Older Mexican Adults” (COSFOMA) a study involving 1,252 adults ≥60 years of age affiliated to the Instituto Mexicano del Seguro Social (IMSS), residents of Mexico City. The IMSS is part of the social protection system in health in Mexico that provides services to salaried workers and their families including medical services and economic benefits such as disability pensions, or paid retirement. In Mexico City, IMSS has 48 primary care medical units attending 36.5% of the population and approximately 50.9% of older adults (7).

A total of 202 individuals from the 1,252 involved in COSFOMA, agreed to participate in the study. In the COSFOMA study, participants were chosen based on their residence through a simple random selection from the list of older adults affiliated to IMSS from the 48 first-level clinics of Mexico City. Written letters delivered to their home addresses were used as invitations to inform them of the nature of the study and to provide them with an appointment at their clinic. When the older adults did not attend the appointment, a phone invitation was made and in some cases a home visit (7). Blood samples were obtained by venipuncture of the median cubital vein using the vacutainer system. Samples collected from April to September 2015 were used for this study.

### Ethics Statement

This research protocol was conducted with the approval of the National Committee of Scientific Research as well as by the Ethics Committee for Health Research of the Instituto Mexicano del Seguro Social (R-2012-785-067). All participants gave written informed consent. This study was performed according to The Code of Ethics of the World Medical Association Declaration of Helsinki.

### Evaluation of reactive oxygen species (ROS)

To evaluate the formation of ROS by fluorometry, 5 μl of serum and 85 μl of PBS 1X buffer was added to a 10 μl solution of dichlorofluorescein diacetate (CAS Number 4091-99-0, Sigma-Aldrich), incubated in darkness for 30 minutes at 37°C for the oxidation of the dichlorofluorescein (DCFH) to the fluorescent compound 2-7-dichlorofluorescin by the presence of peroxide hydrogen to occur. The samples were then read at 498 nm excitation and 522 nm emission (Cytation 5 Cell Imaging Multi-Mode Reader), previously calibrated with a standard curve. Data is shown as pmol/μL of serum.

### Determination of lipid peroxidation

To evaluate lipid peroxidation we measured malondialdehyde (MDA) as a reaction of thiobarbituric acid (CAS Number 504-17-6, Sigma-Aldrich) 50 μl of serum with 25 μl of PBS 1X were added to 50 μl of thiobarbituric acid. The sample was placed in a boiling bath at 94°C for 20 minutes, then centrifuged at 10.500 rpm for 15 min, the supernatant was read at 532 nm with a spectrophotometer (EPOCH), which was previously calibrated with a standard curve. Data is shown as μmol/μL of serum.

### Sample processing and telomere length assessment

Genomic DNA was extracted from peripheral leukocytes by the salting out procedure. Purified DNA samples were aliquoted in a concentration of 10ng/μl and stored at −70 °C until use. For telomere length assessment, we followed the qPCR method published by O’Callaghan and Fenech (21). The number of copies of telomere repeats was determined by the standard curve of Tel STD, while the standard curve of 36B4 STD was used as a housekeeping gene.

After the qPCR reaction on a StepOnePlus Real-Time PCR System (Applied Biosystem), the Ct values of each sample were extrapolated in their corresponding curves by a linear regression test to determine telomere length. The Maxima SYBR Green/ROX qPCR Master Mix 2X (Thermo Scientific, California, USA) was used. The cycling conditions for both genes were: 10 minutes at 95°C, followed by 40 cycles of 95°C for 15 seconds, 60°C for 1 minute, followed by a melting curve. Data is shown as kilobase pairs (kb).

### Frailty

Operationalization of the frailty phenotype was performed using the five criteria proposed by Fried (22). Frail adults are defined as those with three or more of the following criteria: self-report of weight loss, exhaustion, low physical activity, slowness, and weakness (low grip strength). The presence of one or two criteria indicates a pre-frail condition, whereas the absence of criteria indicates a robust or non-frail state (7). Participants were classified as non-frail (score 0), pre-frail (score 1–2), and frail (score 3–5).

### Statistical analysis

In general, the data analysis and visualizations were carried out with free (R software (23) and commercial (SPSS and GraphPad) software. In particular, we used ggplot2 (24) and ggsignif (25) R packages. Quantitative variables are presented as the arithmetic mean and standard deviation (mean ± SD). Qualitative variables are presented as absolute (n) and relative (%) frequencies. BMI was reclassified into nutritional status according to the WHO criteria (underweight <18.5 kg/m^2^, normal weight 18.5–24.9 kg/m^2^, overweight 25–29.9 kg/m^2^, and obesity ≥30 kg/m^2^) to explore the possible correlation between telomere length and nutritional status. In order to explore the distribution of DCFH (pmol/μL of serum), MDA (μmol/μL of serum) and telomere length (kb) between frailty categories (non-frail, pre-frail, and frail), we generated boxplots and calculated their p-value by Mann-Whitney-Wilcoxon test with Bonferroni correction.

The relationship between DCFH and MDA with telomere length by frailty categories was explored by scatter plots and linear regression with Pearson correlation coefficients (r).

The distribution of telomere and oxidative stress among older adult groups was evaluated calculating p-values obtained by Kruskal-Wallis non-parametric test.

The association between three frailty categories with DCFH, MDA, and telomere length was analyzed by multinomial logistic regression analysis with p-values obtained by Student’s t test.

Previous analyses were carried out to determine a possible linear correlation between oxidative stress, lipid peroxidation and telomere length in three conditions: non-frail, pre-frail, and frail old age groups. However, to establish the magnitude of the association against the frailty category, we performed a logistic regression considering the Non-frail group as the reference category to compare the odds of being classified as Pre-frail or Frail considering oxidative stress (DCFH or MDA) or telomere length. The odds ratio (ORs) with 95% confidence intervals (CI 95%) were obtained for precision.

Pearson correlation coefficients, as well as multinomial logistic regression analysis, were adjusted for age, body mass index (BMI), chronic illnesses (0, 1, ≥2), educational level (without studies, 1-6 years, and ≥ 7 years), drinking (binary), smoking (binary), and sex (binary), as possible confounders.

## Results

From the random selection of the COSFOMA study, we invited its 1,252 members and recruited 202 older adults (≥60 years of age) who agreed to participate in this study. The general characteristics as well as reactive oxygen species (ROS) measured by DCFH, lipid peroxidation measured by MDA, and telomere length measurements of the older adults, 133 females (65.8%) and 69 males (34.2%), ranging between 60 and 95 years (mean age: 68.9 ± 7.4) are shown in Table I. Regarding the frailty status, 14.4% of participants were classified as non-frail, 60.4% as pre-frail, and 25.2% as frail.

**Table I.** General characteristics of Mexican older subjects ranging between 60-95 years

| Old age population | Participants | Frequency *n* (%) |
| --- | --- | --- |
| Age (mean ± SD years) | 69.89 ± 7.39 | 202 (100) |
| BMI (mean ± SD kg/m<sup>2</sup>) | 27.71 ± 6.59 | |
| Reactive oxygen species (DCFH) | 50.27 ± 24.58 | |
| Lipid peroxidation (MDA) | 33.53 ± 13.18 | |
| Telomere length (mean ± kb) | 5.31 ± 1.72 | |
| Sex | Female | 133 (65.8) |
|  | Male | 69 (34.2) |
| Nutritional status | Underweight | 8 (3.96) |
|  | Normal weight | 63 (31.19) |
|  | Overweight | 78 (38.61) |
|  | Obesity | 53 (26.24) |
| Education level | Without studies | 16 (7.9) |
|  | 1-6 years | 78 (38.6) |
| | $\geq 7$ years | 108 (53.5) |
| Chronic illnesses | None | 115 (56.9) |
|  | 1 | 57 (28.2) |
| | $\geq 2$ | 30 (14.9) |
| Smoking (currently) | No | 187 (92.6) |
|  | Yes | 15 (7.4) |
| Drinking (currently) | No | 168 (83.2) |
|  | Yes | 34 (16.8) |
| Frailty status | Non-frail | 29 (14.4) |
|  | Pre-frail | 122 (60.4) |
|  | Frail | 51 (25.2) |

The aim of this study was to determine a possible association between oxidative stress measured by reactive oxygen species (DCFH) and lipid peroxidation (MDA), telomere length, and frailty status. We hypothesized that old age subjects with frailty would have increased oxidative stress measurements associated with shorter telomeres when compared with both pre-frail and non-frail subjects.

Contrary to our hypothesis, we observed no significant differences between ROS (DCFH, p= 0.76) or lipid peroxidation (MDA, p= 0.37) among the frailty categories whether divided or not by sex. Between frailty categories, we did find a significant difference with telomere length (p< 0.001). This difference was explained by lower telomere length in the frail group compared against the non-frail (p< 0.001) and the pre-frail (p< 0.001) groups, while the differences between non-frail vs. pre-frail were not statistically significant (p= 0.065) (Figure 1).

**Figure 1.**
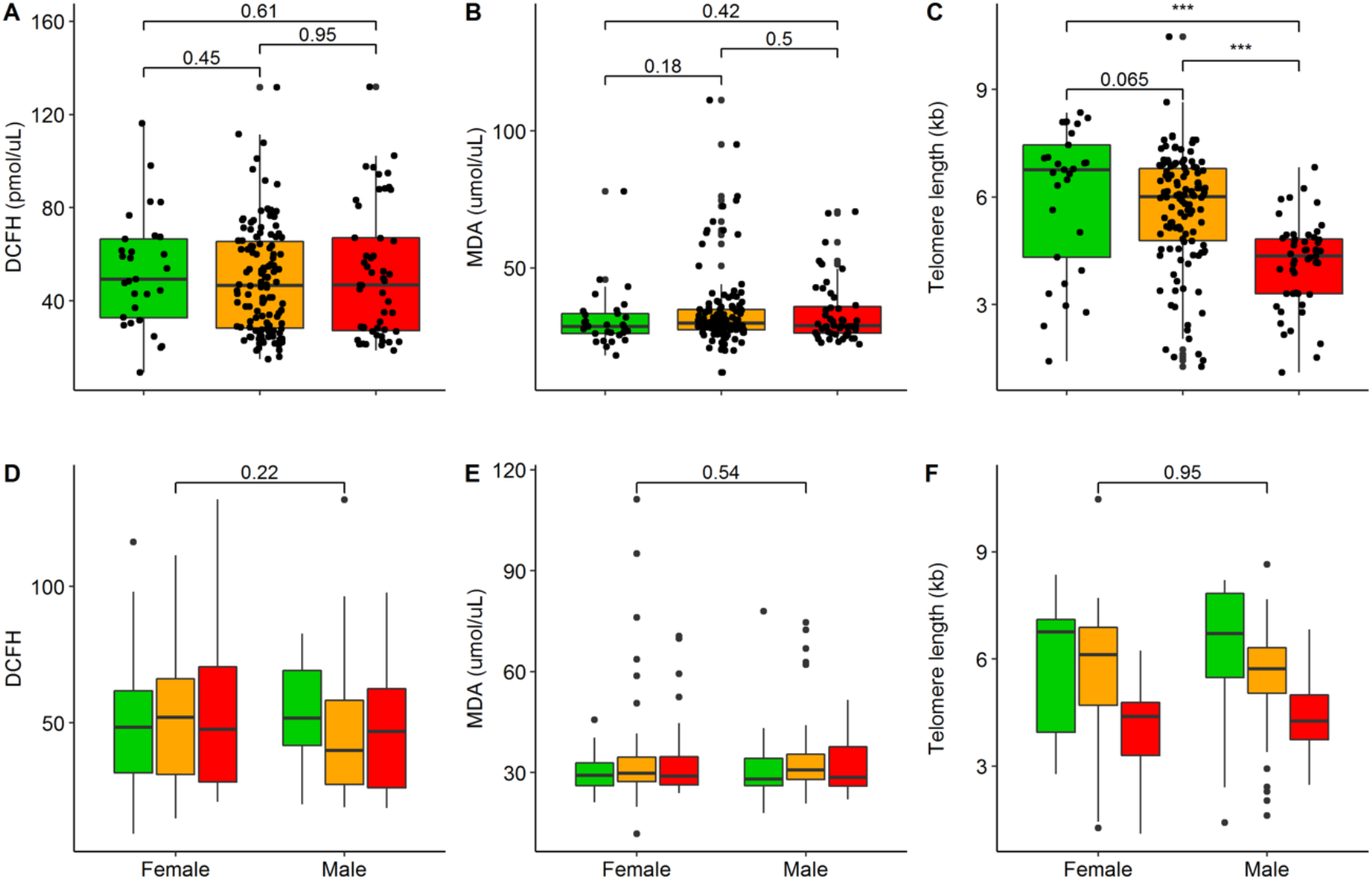
Oxidative stress and telomere length among older adults. This figure shows the distribution of DCFH (p= 0.76), MDA (p= 0.37), and telomere length (p< 0.001), among old age elder groups: Non-frail (green), Pre-frail (orange), and Frail (red). The p-value was obtained by Kruskal-Wallis nonparametric test. The A, B, and C panels show the differences with significant lines within frail categories while panels D, E, and F show the differences with significant lines between females and males by Mann-Whitney-Wilcoxon test. Bonferroni correction < 0.016 was calculated to assign significant differences; p-values < 0.001 (“***”). The p-value was obtained by Kruskal-Wallis nonparametric test.

Next, to assess the possible association between lipid peroxidation (MDA), reactive oxygen species (DCFH) or telomere length with old age groups, we performed a multinomial logistic regression adjusted by age, body mass index, chronic illnesses, educational level, drinking, smoking, and sex. Figure 2 shows the magnitude and precision of the association of oxidative stress and telomere length on pre-frail and frail older adults when considering the non-frail subjects as the reference category. We observed that the odds of being classified in the pre-frail group increases 0.989 per each pmol of DCFH (CI 95% [0.97-1.007] p=0.232), and 1.011 per each μmol of MDA (CI 95% [0.971-1.052] p=0.611), while every kb of telomere (a long telomere) protects 1.33 times from pre-frail classification (OR= 0.749, CI 95% [0.543-1.032] p=0.012). On the other hand, the odds of being classified in the frail group increase 0.986 per each pmol of DCFH (CI 95% [0.964-1.008] p=0.217), and 1.018 per each μmol of MDA (CI 95% [0.971-1.066] p=0.466), while every kb of telomere (a longer telomere) protects 2.28 times from frailty classification (OR= 0.438, CI 95% [0.299-0.641] p<0.001).

**Figure 2.**
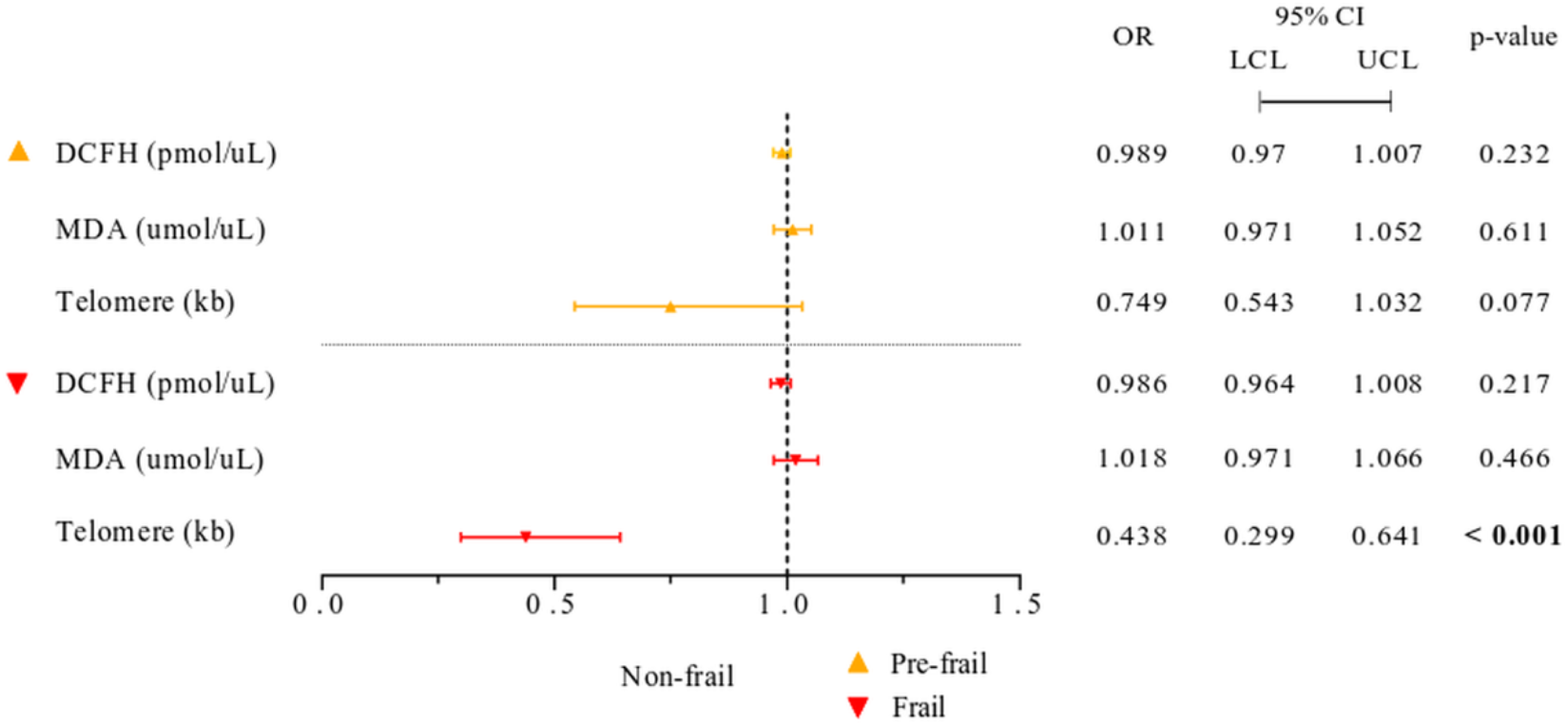
Association of oxidative stress and telomere length in Mexican older adults. The forest plot shows the multinomial logistic regression considering the Non-frail group as the reference category to calculate the odds of being classified as Pre-frail or Frail considering oxidative stress (DCFH or MDA) or telomere length. The odds ratios (OR) with 95% confidence intervals (CI 95%) were obtained for precision and adjusted by age, BMI, educational level, chronic illnesses, smoking, drinking, and sex. Bold text indicates a statistically significant difference (p-value cut-off <0.05).

Finally, we evaluated the relationship between DCFH or MDA on telomere length according to frailty status. First, we calculated the global effect of both DCFH (r = -0.031, p= 0.66) or MDA (r= 0.095, p= 0.18) in telomere length and observed no significant correlation between ROS or lipid peroxidation and telomere length. Then, we performed the analysis for each frailty category (Figure 3).

**Figure 3.**
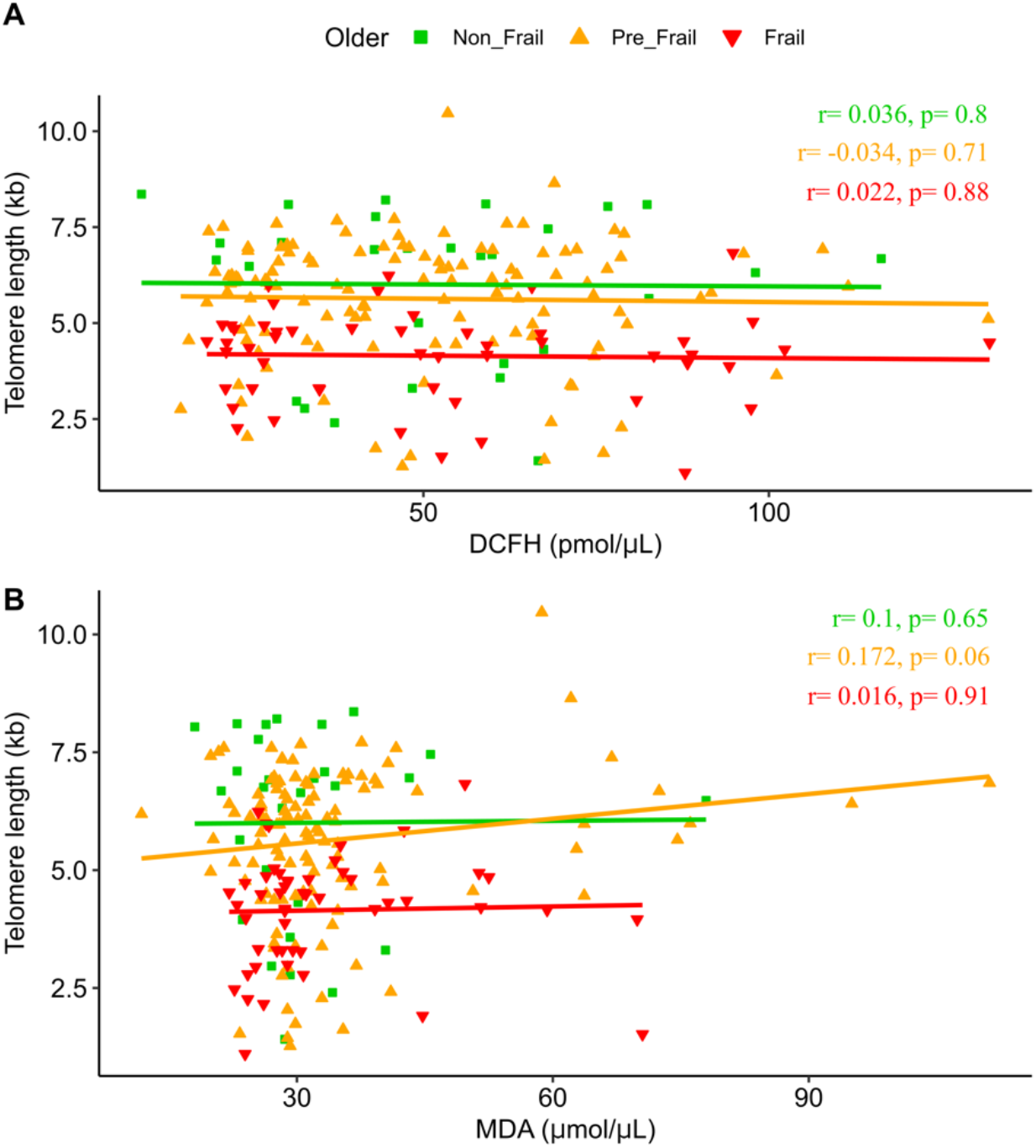
Relationship between oxidative stress and telomere length among frailty categories. The figure shows the correlation analysis for (A) DCFH and (B) MDA, with telomere length. The analysis was carried on the overall older population. Linear regression for non-frail (green; square), pre-frail (orange; triangle), and frail (red; inverted triangle) participants with Pearson correlation coefficients (r) and p-values adjusted by age, BMI, educational level, chronic illnesses, smoking, drinking, and sex are shown.

In addition, as the effect of obesity or adiposity on telomere length has been reported previously (26, 27), and although a negative association of BMI with telomere length is reduced among older people (27), we explored the possible correlation between telomere length and nutritional status (underweight, normal weight, overweight, and obesity), but no significant association was found for either group: underweight (r= 0.640, p= 0.088), normal weight (r= 0.089, p= 0.49), overweight (r= 0.110, p= 0.33), and obesity (r= 0.027, p= 0.85) nor when divided by frailty categories (data not shown).

## Discussion

Frailty, telomere length, and oxidative stress were evaluated in a group of older adults in order to define their association and correlation. A first analysis was carried out to determine our hypothesized distinct distributions between oxidative stress, lipid peroxidation, and telomere length according to frailty groups; however, we observed a significantly different distribution only for telomere length.

Telomere length was found to be associated with pre-frail and frail categories as previously reported (12). This specific variable has become of importance for older adults since, despite the philosophical debate about the definition of frailty, it has shown to have a predictive value on poor health outcomes. Here, we have found that a shorter telomere, which by itself has been previously related to a shorter life span, is associated with the frailty status. Even more, when considering non-frail older adults telomere length as the reference category, short telomeres are associated with pre-frail older adults while even shorter telomere length is associated with frail status. As these results are adjusted by age, body mass index, chronic illnesses, educational level, drinking, smoking, and sex (possible confounders included in our analysis), we consider that telomere length is a strong competitor as a biomarker of frailty status. To achieve this, this study must be replicated in other populations in order for telomere length to be considered as a biomarker of frailty since telomere length is affected by environmental factors (28).

Elseways, telomere length has been found to have modifiable factors such as smoking and obesity, which are associated with shorter telomeres while exercise and a healthy diet preserve telomere length (4, 29). Furthermore, a recent study showed that frailty can be modified by exercise and nutrition, reversing the frailty state to a non-frail (30). Hence, telomere length determination as a part of frailty diagnosis could help apply prompt intervention, either in pre-frail or frail status, on the above mentioned factors in order to reverse this condition. If pre-frail status and frailty are assessed on time, more costly medical outcomes could be prevented and quality of life improved. Research in this direction may help determine frailty in a more quantitative manner with a biological background. Although telomere length determination is currently not widely available, its future implementation could be considered as part of the frailty diagnosis, as a non-invasive tool.

On that matter, the association of frailty status with telomere length could mean that besides its definition, frailty has a biological counterpart that changes as frailty status does. Hence, we evaluated oxidative stress as a probable cause for telomere shortening that could be assessed as telomere length and found a mild effect of MDA on telomere length (r=0.17; p=0.06) for the pre-frail category. MDA is a marker for lipid peroxidation, oxidative degradation of lipids that follow the excess of free radical formation. The increase of MDA in pre-fail patients and the decrease in telomere length between pre-frail and frail patients suggests that oxidative stress has an effect on telomere length regarding frailty status in older adults.

The main limitation of the present study is that even when the cohort used is constituted by 1,252 individuals, we were only able to include 202 participants, meaning our recruitment skills were insufficient. The hypothesis generated regarding the present data could be proven with a longitudinal study that would provide further information regarding the effect of MDA on telomere length and frail status. Also, a more thorough analysis of oxidative stress, specifically the measurement of 8-hydroxy-2’-deoxyguanosine, a specific biomarker of oxidative stress in DNA (31), would enrich the knowledge on the effect of these two biological outcomes of frailty.

It is worth mentioning that when not considering drinking, smoking, and chronic illnesses as confounder variables, we observed a linear association between MDA and telomere length (p= 0.089). We are convinced that an ideal approach would have these as exclusion criteria in order to avoid masking this biological measurement.

In conclusion, we reaffirmed an association of telomere length in the frailty phenotype and found a low association between lipid peroxidation and telomere length. Whether oxidative stress is the main cause of telomere length shortening in frail old adults remains to be proven. Moreover, we found no effect of drinking or smoking as confounder variables regarding telomere length or oxidative stress in frail old adults for this study.

## Abbreviations

DM: diabetes mellitus
ROS: reactive oxygen species
COSFOMA: Cohort of obesity, sarcopenia, and frailty of older Mexican adults
IMSS: Instituto Mexicano del Seguro Social
DCFH: dichloro-dihydro-fluorescein
MDA: malondialdehyde
BMI: body mass index

## Acknowledgements

The authors thank Mario Enrique Rendón-Macías for his comments on the data analysis and visualization. This study was performed in partial fulfillment of the requirements for the PhD degree of JDME who thanks the Posgrado en Ciencias Biológicas, Biología Experimental, UNAM.

